# Skeletal muscle mitochondria contain nuclear-encoded RNA species prior to and following adaptation to exercise training in rats

**DOI:** 10.1101/2025.01.12.631800

**Authors:** Jessica L. Silver, Séverine Lamon, Stella Loke, Gisella Mazzarino, Larry Croft, Mark Ziemann, Glenn. D. Wadley, Adam J. Trewin

## Abstract

Skeletal muscle mitochondria adaptation to exercise training is mediated by molecular factors that are not fully understood. Mitochondria import over 1000 proteins encoded by the nuclear genome, but the RNA population resident within the organelle is generally thought to be exclusively encoded by the mitochondrial genome. However, recent *in vitro* evidence suggests that specific nuclear-encoded miRNAs and other non-coding RNAs (ncRNAs) can reside within the mitochondrial matrix. Whether these are present in mitochondria of skeletal muscle tissue, and if this affected by endurance training - a potent metabolic stimulus for mitochondrial adaptation - remains unknown. Rats underwent four weeks of moderate intensity treadmill exercise training, then humanely killed and tissues collected for molecular profiling. Mitochondria from gastrocnemius skeletal muscle were isolated by immunoprecipitation, further purified, then the resident RNA was sequenced to assess the mitochondrial transcriptome. Exercise training elicited typical transcriptomic responses and functional adaptations in skeletal muscle, including increased mitochondrial respiratory capacity. We identified 24 nuclear-encoded, coding or non-coding RNAs in purified mitochondria, in addition to 50 nuclear-encoded miRNAs that met a specified abundance threshold. Although none were differentially expressed in the exercise vs control group at FDR<0.05, exploratory analyses suggested that the abundance of 3 miRNAs were altered (*p*<0.05) in mitochondria isolated from trained compared with sedentary skeletal muscle. We report the presence of a specific population of nuclear-encoded RNAs in the mitochondria isolated from rat skeletal muscle tissue, which could play a role in regulating exercise adaptations and mitochondrial biology.

**KEY POINTS:**

- Mitochondria are a central metabolic and signalling hub, particularly in tissues with high energy demand such as skeletal muscle. Their ability to adapt to stimuli such as exercise training is directly related to improved metabolic health. Understanding the factors that regulate these adaptive processes is therefore essential.
- Mitochondria contain RNA encoded by their own genome (mtDNA); typically are not thought to contain nuclear-encoded RNA.
- Here, we report that mitochondria from rat skeletal muscle do indeed contain a select population of nuclear-encoded protein coding and non-coding RNAs, including 3 microRNAs whose expression tended to be altered by exercise training.
- These findings suggest that mitochondria-localised nuclear-encoded RNAs may play a role in mediating beneficial adaptive responses to exercise.

## INTRODUCTION

Exercise training elicits positive, systemic adaptations within the cardiovascular, endocrine, and muscular systems, which reduce the risk of chronic metabolic conditions such as type 2 diabetes and cardiovascular disease (1). Within skeletal muscle, endurance exercise training improves the capacity to oxidise carbohydrate and lipid substrates, increases lactate threshold and can delay the onset of fatigue (1, 2). These intramuscular adaptations are reflected by an increase in skeletal muscle mitochondrial content (3). Mitochondria are a central metabolic hub for the integration of fundamental cellular processes. The abundance and function of mitochondria within a cell are related to its ability to maintain bioenergetic, ion and redox homeostasis (4). Through the process of mitochondrial biogenesis, repeated bouts of endurance exercise increase the abundance of mitochondrial constituents and functional capacity, which are hallmarks of a shift towards a more oxidative muscle fibre type and results in improved endurance exercise capacity (2, 3, 5). However, the complex molecular events and transcriptional networks governing mitochondrial adaptations at the whole muscle level remain incompletely understood, and even less is known regarding the regulation of exercise-induced adaptations within mitochondria.

One means of transcriptional and translational regulation involves non-coding RNA (ncRNA) MiRNAs are a class of short non-coding RNAs (typically ∼22 nt) that selectively recognise untranslated regions of a target mRNA (6) and most commonly repress translation by degrading a target mRNA and thus decreasing translational efficiency (7). All known miRNAs are transcribed from the nuclear genome (8), before being processed into mature miRNAs that are incorporated into RNA-induced silencing complexes (RISC) usually within the cytoplasm (9). Endurance training induces changes in skeletal muscle miRNA expression in both human (1, 10, 11) and animal (12, 13) models at the whole tissue level. Long non-coding RNAs (lncRNAs) are transcripts >200 nt in length that can function in multiple ways, including by binding to complementary sequences in regulatory elements of DNA to enhance or repress transcription of mRNA, by binding to mRNAs to regulate protein translation, or via physical interactions with proteins to alter their function (14). We and others have demonstrated that various lncRNAs play a role in exercise adaptations in myocytes (15–19). However, only a small fraction of currently annotated miRNA and lncRNA transcripts have been investigated experimentally to determine their biological or molecular function(s) (20).

Mitochondria retain a conserved circular genome (mtDNA) containing 37 genes, 13 of which encode mRNAs that are translated into essential protein subunits of the electron transport system necessary for oxidative phosphorylation (21, 22). In addition, over 1,000 nuclear-encoded genes are required for various aspects of mitochondrial function, but these are translated in the cytosol then imported into mitochondria as nascent protein via the TIM/TOM complex (23). While there is no equivalent mechanism for the bulk uptake of nuclear-encoded RNA into mitochondria, certain nuclear-encoded RNA species have been shown to be localised within mitochondria. For instance, specific small RNAs, such as miRNAs can localise within mitochondria of rat heart (24), mouse heart (25–27) and human myoblasts, kidney and cancer cells (28–31), among others (reviewed in (32)). There is also emerging evidence from *in vitro* studies that a few specific nuclear-encoded ‘long’ RNAs are imported into mitochondria, such as 5S rRNA, and lncRNAs *RPPH1, RMRP* and *mitolnc* (22, 33–35), although this has not yet been well characterised (36). Given the potential regulatory effects that nuclear-encoded RNA could elicit particularly within mitochondria of metabolically active tissues, it is important to determine which ones are mitochondria-localised. Despite this, an unbiased whole-transcriptomic screen for both small (e.g. miRNAs) and long RNA species (i.e. mRNA and lncRNAs) has not previously been performed on mitochondria from skeletal muscle. The aim of this study was to characterise the whole transcriptome contained within mitochondria isolated from rat skeletal muscle, and to investigate whether this is altered in association with mitochondrial adaptations to exercise training.

## RESULTS

### Physiological responses to exercise training

Rats performed moderate intensity treadmill exercise training (ExT) or remained sedentary (Sed) for 4 weeks and were assessed for the ensuing physiological and molecular adaptations in the heart and skeletal muscle. Body weight was not different between ExT and Sed rats after the 4-week exercise training intervention (Figure 1A). Consistent with the known beneficial effects of exercise training that induce physiological cardiac hypertrophy (37), 4-week exercise training led to greater whole heart mass in ExT compared to Sed (*p*=0.038; Figure 1B), although left ventricle free wall mass was not significantly greater (*p*=0.12; Figure 1C).

**Figure 1:**
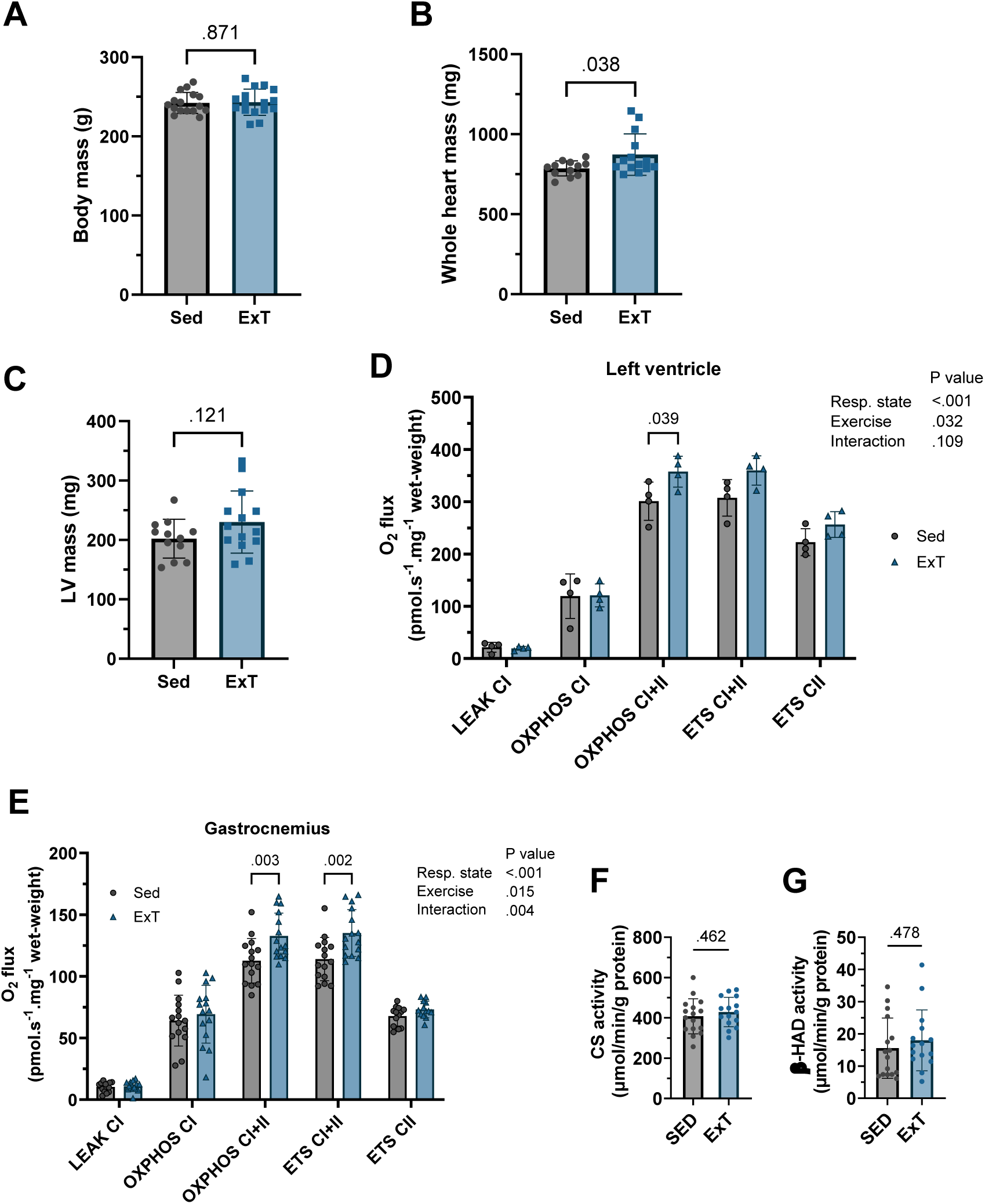
Effects of 4 weeks of treadmill exercise training (ExT) or sedentary control (Sed) on rat whole-body characteristics, and heart and skeletal muscle mitochondrial respiratory capacity. Rat body mass A) at 9 weeks of age after 4 week treadmill exercise training (ExT) or sedentary control (Sed); data are mean(SD) analysed by t-test for *n*=16/grp. B) Heart weight, and C) left ventricle outer wall mass (*n*=12 Sed, *n*=14 ExT; tissue mass data for 6 animals not recorded due to equipment fault). D) Left ventricle and E) red portion of gastrocnemius mitochondrial oxygen consumption per mg tissue during non-ADP stimulated state-4 (Leak) respiratory state, 2.5 mM ADP-stimulated oxidative phosphorylation (OXPHOS) or uncoupled electron transport system capacity (ETS) supported by complex I and/or II substrates. Left ventricle respiration analyses was performed on a subset of animals (*n*=4 per group), gastrocnemius tissue analysis was performed on *n*=15 per group. Data are mean(SD) analysed by two-way ANOVA (left ventricle) or Mixed-effects analysis (Gastrocnemius) with repeated measures, with Bonferroni post-hoc test. F) Citrate synthase and G) 3-hydroxyacyl-CoA dehydrogenase (β-HAD) maximal enzyme activity from red portion of the gastrocnemius; data are mean(SD) analysed by t-test for *n*=16 (Sed) and *n*=15 (ExT).

### Mitochondrial functional adaptations after exercise training

Mitochondrial respiratory function was assessed in heart and skeletal muscle. In left ventricle tissue from a subset of animals (*n*=4 per group), mitochondrial respiratory capacity (O_2_ consumption) measured by high resolution respirometry was greater during ADP-stimulated oxidative phosphorylation (OXPHOS) supported by convergent complex I+II substrates (malate/glutamate + succinate) in ExT compared with Sed (Figure 1D). Similarly, in the red portion of the gastrocnemius skeletal muscle (*n*=15 per group), ExT led to increased rates of mitochondrial respiration under OXPHOS CI+II, as well as uncoupled electron transport system maximal capacity CI+II supported respiration (ETS CI+II), which were both ∼18% greater with ExT compared to Sed (Figure 1E). These substrate-specific effects of ExT on mitochondria occurred independently of any changes in maximal citrate synthase or β-HAD activities in gastrocnemius skeletal muscle (Figure 1F & 1G). Taken together, these data indicate that 4 weeks of moderate intensity treadmill exercise induced expected mitochondrial adaptations in rat cardiac and skeletal muscle.

### Transcriptomic responses to exercise training in skeletal muscle and heart tissue

To understand the molecular pathways underlying these adaptations, we performed whole-transcriptome RNA sequencing on cardiac and skeletal muscle whole tissue (Supp Fig 1A&B). In heart tissue (outer wall of the left ventricle) from a subset of animals (*n*=7 per group), there were no genes considered to be differentially expressed (DE) with a strict false discovery rate threshold (FDR<0.05; Supplementary Dataset 1). However, for exploratory purposes, there were 121 genes differentially expressed with an individual p-value less than 0.01 (Figure 2A; Supplementary Dataset 1). Consistent with the increase in LV mitochondrial respiratory capacity, pathway enrichment analysis revealed that various mitochondria-related GO Biological Processes terms were enriched including ‘*Mitochondrial translation*’, ‘*Mitochondrial gene expression*’, and ‘*Aerobic respiration*’ (Figure 2B), despite individual OXPHOS subunit genes not being significantly DE at the FDR 0.05 level (Figure 2C).

**Figure 2:**
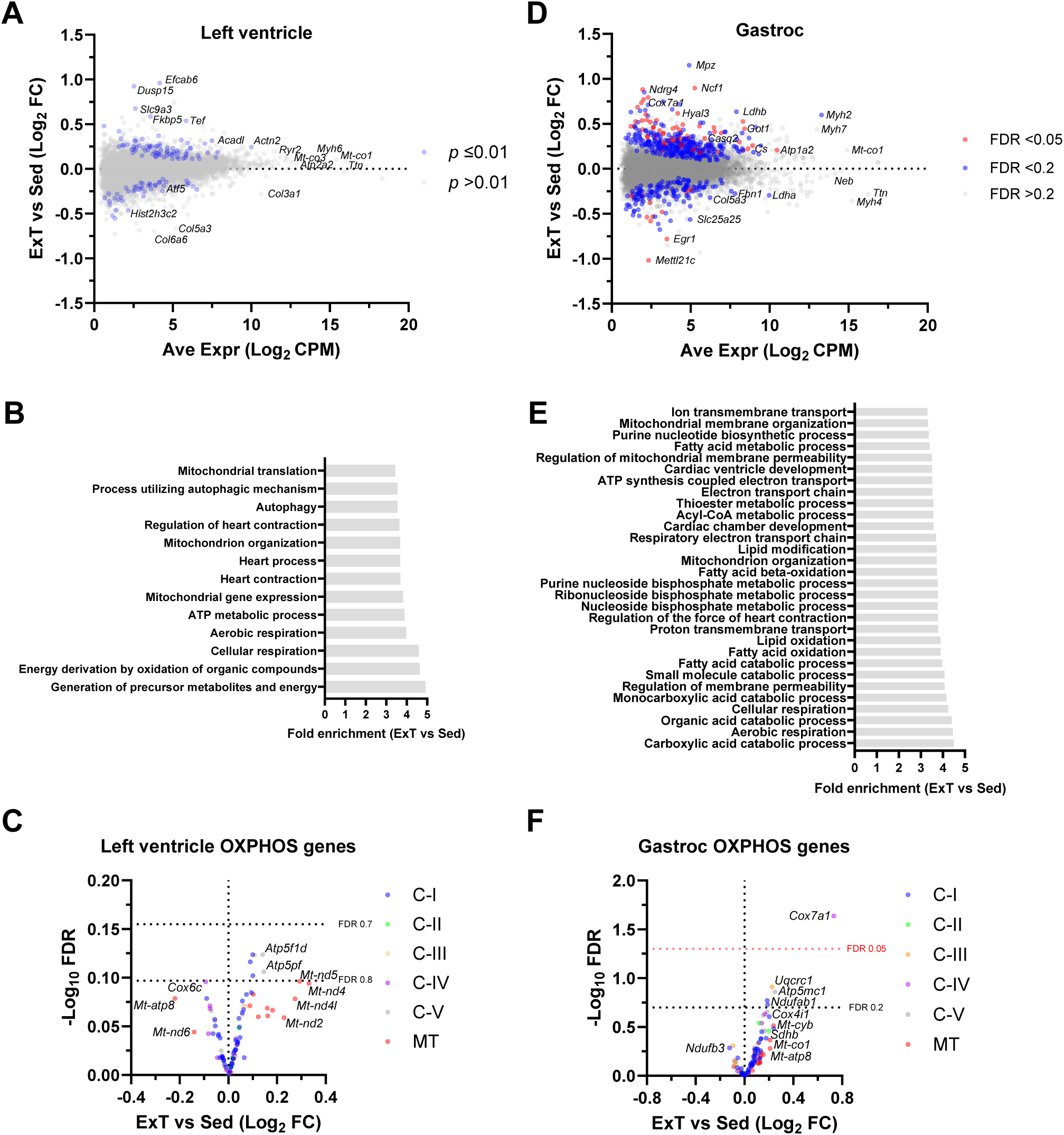
Transcriptome of rat left ventricle and gastrocnemius skeletal muscle tissue after 4 weeks of treadmill exercise training (ExT) or sedentary control (Sed). Whole tissue lysate from left ventricle and the red portion of the gastrocnemius was subjected to total RNA sequencing. **A)** Left ventricle MA plot of all transcripts (≥1 CPM in all samples), positive fold-change (log2FC) indicates upregulation in ExT at unadjusted *p*<0.01 (dark grey) and *p*>0.01 (light grey) for *n*=7 samples per group. **B)** GO Biological Process terms enriched in left ventricle with ExT vs Sed; pathway FDR<0.2. **C)** Volcano plot of nuclear-encoded genes for subunits of OXPHOS complexes I-V (CI – CV) as well as mtDNA-encoded genes (MT) in left ventricle for ExT vs Sed. **D)** Gastroc MA plot of all transcripts (≥1 CPM in all samples), positive fold-change (FC) indicates upregulation in ExT at FDR<0.05 (red) and FDR<0.2 (blue) for *n*=15 samples per group. **E)** GO Biological Process terms enriched in gastroc with ExT vs Sed; pathway FDR<0.1. **F)** Volcano plot of nuclear-encoded genes for subunits of OXPHOS complexes I-V (CI – CV) as well as mtDNA-encoded genes (MT) in gastroc for ExT vs Sed.

In the type-I fibre-rich portion of gastrocnemius skeletal muscle tissue, there were 30 upregulated and 4 downregulated genes (FDR<0.05) in the ExT group compared with Sed (Figure 2D, Supplementary Dataset 2). At a less stringent FDR of <0.2 chosen for the purpose of discovery, we observed 347 upregulated and 140 downregulated genes in the ExT group compared with Sed including 7 long non-coding RNAs and 2 pseudogenes in the exercise trained state (Supplementary Dataset 2). Pathway analyses revealed that several key mitochondrial and metabolic processes were upregulated in response to ExT, including ‘*Mitochondrial membrane organization*’, ‘*Fatty acid metabolic process*’, and ‘*Aerobic respiration*’ (Figure 2E). At the individual gene level, *Cox7a1* was significantly upregulated in response to ExT at FDR <0.05 and other OXPHOS genes tended to increase (FDR 0.05-0.2; Figure 2F). Finally, our transcriptome data also provides evidence of the expected slow muscle fibre-type shift in response to exercise training. Specifically, gene expression of isoforms of myosin light and heavy chains as well as troponins associated with fast-twitch fibres (38) were decreased in ExT compared with Sed (p<0.05; Supplemental Figure 1C-F).

### Enrichment of nuclear-encoded RNAs within isolated mitochondria of skeletal muscle

RNA contained within mitochondria is generally thought to be exclusively the transcriptional product of mtDNA, which encodes 13 essential OXPHOS subunits along with ribosomal and transfer RNAs. To investigate whether nuclear-encoded transcripts are also present within the mitochondrial matrix and/or intermembrane space of skeletal muscle tissue, we isolated mitochondria from gastrocnemius skeletal muscle by immunoprecipitation of a protein localised at the outer mitochondrial membrane (TOMM20). These isolated mitochondria were then enzymatically purified using RNase-A to remove RNAs bound to ribosomes on the outer membrane of the intact mitochondrial organelle (Figure 3A), yielding a highly-purified mitochondrial fraction as described by our group (39). The total mitochondrial RNA population contained within the intact organelle was then collected and used to construct two separate library types for RNA-sequencing: firstly, a ribosomal RNA depleted ‘total’ RNA library to assess the protein coding and non-coding transcriptome; and secondly, a small RNA library (transcripts <∼200 nt) to identify microRNAs. Using this approach, we generated total RNA and small RNA libraries from isolated and purified mitochondria which generated on average 27 and 33 million reads per mitochondrial library, respectively.

**Figure 3:**
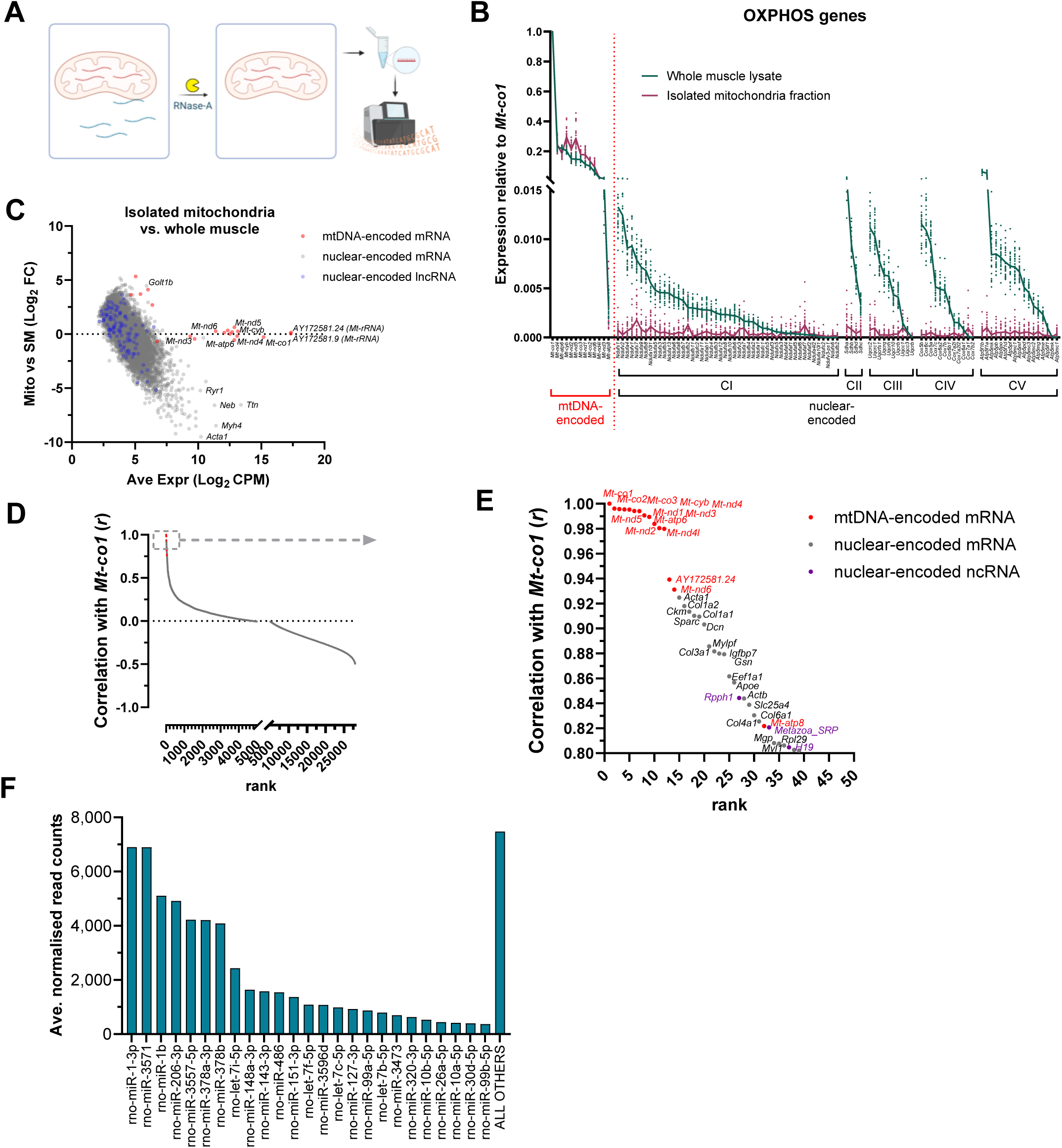
Transcriptome of isolated mitochondria from red portion of rat gastrocnemius skeletal muscle. **A)** Experimental overview of mitochondrial isolation from gastrocnemius muscle tissue. Isolated mitochondria were enzymatically purified with RNase-A to remove transcripts on the outer mitochondrial membrane. RNA extracted from these purified mitochondria and their respective whole tissue lysate then underwent RNA-sequencing. **B)** Purity of the isolated mitochondria preparation indicated by the relative absence of nuclear-encoded transcripts for subunits of OXPHOS complexes I to V in isolated mitochondria fraction compared with whole muscle lysate samples. Data are RNA-seq counts for each gene expressed relative to *Mt-co1* (the most abundant mRNA in the mitochondrial fraction), presented as mean and individual data points for *n*=30 animals (Sed and ExT combined). **C)** MA-plot of isolated mitochondria (Mt) relative to whole skeletal muscle (SM); mtDNA-encoded transcripts are shown in red, all nuclear-encoded transcripts in grey. **D)** Correlation of all genes (>1 CPM) with *Mt-co1* in isolated mitochondria (*n*=30 animals, Sed and ExT combined); red and grey points represent mtDNA-encoded and nuclear-encoded transcripts, respectively. Positive correlation indicates mitochondrial localisation, negative correlation suggests non-mitochondrial localisation. **E)** Enlarged plot of panel D, showing transcripts with a correlation greater than *r*=0.8 with *Mt-co1*. **F)** The top 25 most highly expressed miRNAs in rat skeletal muscle mitochondria. Data are the average normalised read counts per miR across all samples (*n*=25 animals, Sed and ExT combined).

First, to confirm the purity of the mitochondria sample, we took advantage of the fact that mtDNA-encoded OXPHOS transcripts are detectable in both whole tissue and isolated mitochondrial fractions, whereas nuclear-encoded OXPHOS transcripts are not expected to be detected in isolated mitochondria. Indeed, isolated mitochondria contained all 13 mtDNA-encoded mRNAs at approximately equal abundance as their respective whole muscle tissue sample when normalised to *Mt-co1* as an internal control, and were essentially devoid of all nuclear-encoded OXPHOS transcripts (*n*=30 ExT & Sed combined, Figure 3B), indicating a high degree of purity. Furthermore, several of the most highly abundant mRNAs in skeletal muscle tissue, such as *Ryr1*, *Neb*, *Ttn*, *Myh4* and *Acta1* were scarcely detected in the mitochondrial fraction (i.e., ∼40 to 700-fold depletion; Figure 3C). Collectively, this was entirely consistent with our validation studies using highly purified mitoplasts (39). Intriguingly, we detected nuclear-encoded lncRNAs and mRNAs in the mitochondrial samples, albeit with relatively low levels of abundance relative to the mtDNA-encoded mRNAs (Figure 3C). Nonetheless, these included a few specific transcripts that appeared to be enriched in the mitochondrial fraction such as *Golt1b* (Figure 3C and Supplementary Dataset 3).

To identify *bona fide* mitochondria-localised RNA transcripts, we devised an analysis strategy which exploits the minor between-sample variability in completeness of the RNase-A digestion process. Across all mitochondrial samples (*n*=30, ExT & Sed combined), we correlated the counts of every gene with those of *Mt-co1*, because it is *a*) the most highly abundant mtDNA-encoded mRNA, and *b*) unaffected by RNase-A due to its mitochondrial origin. Therefore, positive correlation with *Mt-co1* will reveal mitochondria-localised transcripts (i.e. all mtDNA-encoded transcripts) since these would all be unaffected by the RNase treatment in intact organelles. Conversely, non-mitochondrial localised transcripts would negatively correlate with *Mt-co1* because a more ‘complete’ RNase treatment will preferentially decrease their abundance in that sample. Indeed, using this approach, we confirmed the expected strong positive correlation between *Mt-co1* and all other mtDNA-encoded genes, whereas the vast majority (>99%) of nuclear-encoded genes (which would be expected to be localised to the cytosol or nucleus) had either a negative or no more than a weak positive (*r*<0.4) correlation with *Mt-co1* (Figure 3D). Based on this robust approach of assessing the mitochondrial localisation of transcripts, we identified 24 nuclear-encoded transcripts that exhibited a strong positive correlation (*r*>0.8) with *Mt-co1* (Figure 3E). Notably, this included the mitochondrial localisation of *Rpph1*, a nuclear-encoded RNA component H1 of RNase-P involved in processing of tRNAs which has previously been experimentally demonstrated to be imported into mitochondria (34, 40, 41). We detected two other non-coding RNAs which displayed mitochondrial localisation, *H19* and *Metazoa SRP*. We also identified nuclear-encoded mRNAs including *Acta1*, along with transcripts that encode isoforms of collagen proteins (including *Col1a2*, *Col1a1*, and *Col3a1*) which had strong correlations (*r*>0.8) with *Mt-co1*, suggesting that a small pool of these mRNAs are mitochondria-localised (Figure 3E and Supp Fig 2). Collectively, these data suggest that a small number of specific RNAs may be imported into mitochondria in skeletal muscle and could play as-yet unknown role(s) in mitochondrial biology.

### Enrichment of miRNAs within isolated mitochondria

We next assessed whether a population of miRNAs reside within isolated mitochondria from skeletal muscle. To do this, we sequenced small RNA libraries constructed from RNA from isolated mitochondria from the red gastrocnemius. Approximately 70% of reads sequenced from small RNA libraries mapped to the rat miRbase transcriptome across all libraries (Sed + ExT combined, *n*=25). Less than 1% (sedentary = 0.24±0.16%, trained = 0.30±0.15%) of total reads mapped to known rat miRNA sequences (Supp Fig 2E). 764 mature miRNAs have been identified within the rat genome and are annotated in miRbase (42) yet only small number of these were detected in mitochondria isolated from gastrocnemius tissue (Supp Fig 2F). After applying thresholds to include miRNAs with >100 normalised read counts, we detected 50 mature miRNAs in mitochondria, with miR-1-3p being the most abundant (Figure 3F).

### Effects of exercise training on the transcriptome of isolated mitochondria

Having identified mitochondria-localised nuclear-encoded RNAs and given that exercise training improved skeletal muscle mitochondrial function and altered gene expression, we next investigated if the subcellular transcriptome of the mitochondrial organelle is altered by exercise training. Owing to the inherently greater between-sample variability of a subcellular fraction compared to whole tissue lysate, there were no genes that were considered to be differentially expressed between ExT and Sed within the total RNA transcriptome of isolated mitochondria at the strict FDR <0.05 threshold (Figure 4A). However, there were 365 genes with an individual *p*-value <0.05 (Supplementary Dataset 4A). None of the RNAs that were positively correlated with *Mt-co1* (shown in Fig 3E) were among those affected by ExT compared with Sed at *p*<0.05 in (Figure 4A).

**Figure 4:**
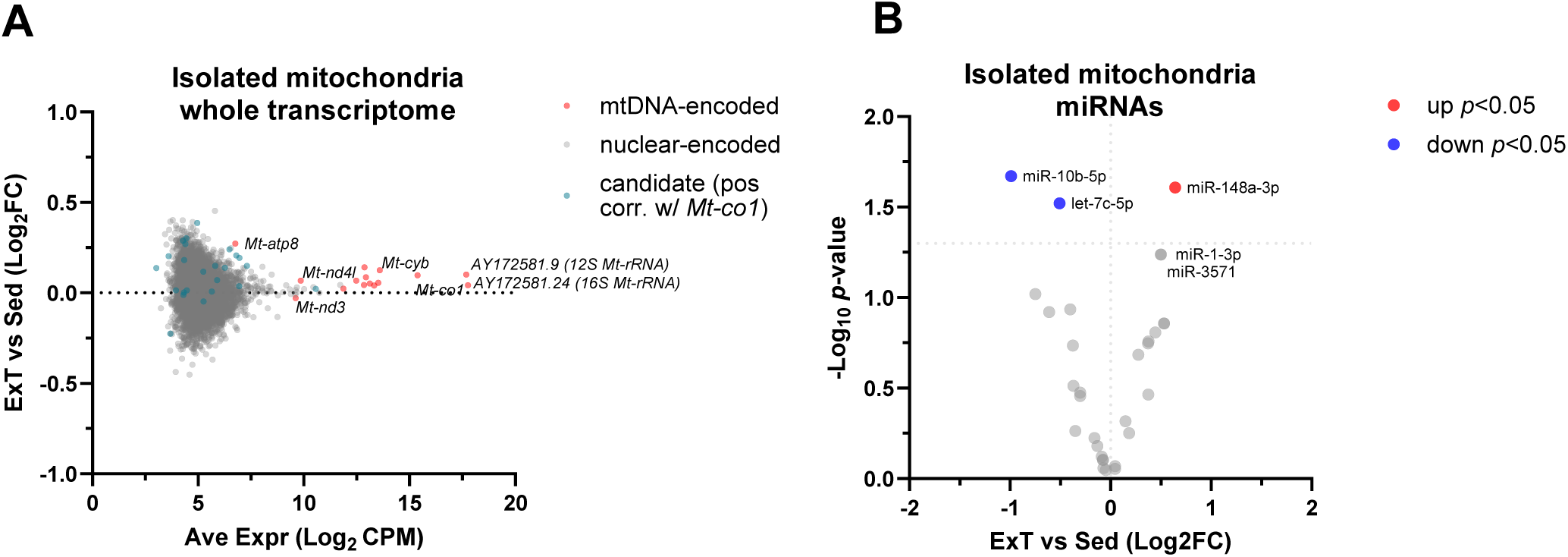
Effects of 4 weeks exercise training on the transcriptome of mitochondria isolated from rat gastrocnemius skeletal muscle. **A)** MA plot of all transcripts (≥2 CPM in all samples) for *n*=15 samples per group, no genes were significantly DE between ExT and Sed at FDR<0.05. mtDNA-encoded transcripts are shown in red, nuclear-encoded transcripts in grey, and transcripts identified as being mitochondria-localised candidates from correlation with *Mt-co1* (from Fig 3E) shown in green. **B)** Volcano plot of miRNAs detected in isolated mitochondria (>100 counts in >80% samples) for ExT vs Sed. No miRNAs were differentially expressed at the FDR<0.05 threshold, data shown are *p*-values for *n*=12 (Sed) and *n*=13 (ExT).

Of the miRNAs detected in mitochondria, there was 1 upregulated (miR-148a-3p) and 2 downregulated (miR-10b-5p and let-7c-5p) in ExT compared with Sed at an individual *p*-value <0.05, along with a trend (*p*=0.057) for both miR-1-3p and miR-3571 to be upregulated (Figure 4B, Supplementary Dataset 4B). None of these candidate miRNAs had experimentally validated mRNA targets encoded by mtDNA, however, there were 45 validated nuclear encoded mRNA target interactions reported in previous literature. Two experimentally validated interactions were found with miR-10b-5p, *Bdnf1* and *Rhoc* yet we did not detect these mRNAs in our mitochondrial dataset. The remaining experimentally validated mRNA interactions were with miR-1-3p. Notable mRNA targets of miR-1-3p included *Igf1* (43), involved in cell development and apoptosis, *Mef2a* (44), a transcription factor that mediates calcium-dependant changes in gene expression in skeletal muscle, and the mitochondria-localised isoform of the antioxidant enzyme, *Sod1* (45). Both *Igf1* and *Sod1* mRNA each had weak, yet positive correlations with *Mt-co1* (*r*=0.19, *r*=0.15, respectively), suggesting that some of their mRNA may reside within the mitochondrial compartment, where it could be targeted by miR-1-3p.

## DISCUSSION

The main findings were that mitochondria isolated from rat skeletal muscle samples contained ∼50 mature miRNAs and 24 nuclear-encoded RNA transcripts. Exercise training led to the expected improvement in skeletal muscle mitochondrial function and significantly impacted the overall skeletal muscle transcriptome, along with a few specific miRNAs that were changed within the subcellular mitochondrial transcriptome following exercise training.

It is well established that nuclear-encoded mRNAs localise to ribosomes on the outer mitochondrial membrane for translation and importation into mitochondria as nascent proteins (46), but less is known about RNAs that may translocate across the mitochondrial membranes into the matrix as RNAs. We detected a small number of nuclear-encoded RNA transcripts localised to the mitochondria. One transcript, *Rpph1* is an RNA component of the RNase P ribonucleoprotein complex responsible for processing tRNAs has been experimentally demonstrated to be imported into mitochondria (34, 41, 47), thus validating our finding. Intriguingly, we also identified other nuclear-encoded RNAs not previously known to reside within mitochondria. The lncRNA H19 is an important gene in epigenetic control of early development and growth and has been shown to regulate skeletal muscle fibre type in response to exercise training (18). Another ncRNA, the Metazoan signal recognition particle RNA (*Metazoa SRP*) is a component of the SRP which targets specific proteins to the endoplasmic reticulum. Moreover, a number of nuclear-encoded mRNAs were detected within mitochondria including isoforms of collagen (*Col1a2*, *Col1a1*, and *Col3a1*) and actin *Acta1* and *Actb*. These nuclear-encoded mRNAs would be incompatible for recognition and translation on mitochondrial ribosomes, yet they could regulatory function within the mitochondrial matrix by via interactions directly with mt-RNAs or with RNA-binding proteins. Whether a small population of these RNAs are imported into mitochondria indiscriminately or whether they are imported for specific functional purposes will be important to elucidate. Earlier work by Wang et al. (34) has demonstrated a mechanism in which the PNPase recognises specific stem-loop secondary structures of RNA allowing it to import RNA species containing that structure into mitochondria (34). Their work showed that PNPase was required for the import of RNase P, 5S rRNA, and MRP RNAs (34). Whether this or another similar mechanism could explain our findings will also be important to investigate in future studies.

The localisation of microRNAs in mitochondria from various cell types is better established than other nuclear-encoded RNA species (48). In the present study, less than 1% of total sequenced reads mapped to known mature miRNAs. While this proportion is lower than that reported by others elsewhere (2.6% and 4.2% of total reads mapping miRNAs in HeLa and HEK293 cells, respectively (31)), this may be due in part to a lack of RNase treatment in the aforementioned study, and because only 764 miRNAs have been annotated in the rat genome compared to 1,978 and 2,656 mature miRNAs annotated in the mouse and human genomes, respectively (42). Of the reads mapping mature rat miRNAs, the skeletal-muscle enriched miR-1-3p and miR-206 were among the most highly expressed miRNAs in rat skeletal muscle mitochondria. MiR-1 is well characterised in the literature and has been previously shown to localise in the mitochondria of human primary myoblasts (29), C2C12 myotubes (27) and rat cardiac muscle in vivo (49). Crosslinking immunoprecipitation coupled with deep sequencing further demonstrated that the mitochondria-localised miR-1 directly associated with AGO2 and transcripts encoded by the mitochondrial genome including *Mt-Co1*, *Mt-Nd1*, *Mt-Cyb*, *Mt-Co3* and *Mt-Atp8* (27). Interestingly, miR-1 reportedly increased, rather than inhibited, the translation of mitochondrial transcripts in C2C12 myotubes (27). Given that several miRs showed tendencies to be differentially expressed within mitochondria in response to exercise training, it will be worth considering the functional implications of this in future studies.

In our study, exercise training led to canonical mitochondrial adaptations in the heart and skeletal muscle, both highly metabolic tissues. Our analyses were conducted on these tissues which were collected 24 h after the final session of exercise. Therefore, we characterised the transcriptome of the chronically trained, yet rested state. It is possible that tissue collection at an earlier post-exercise time point would capture the distinctive transcriptional responses that occur following an acute bout of exercise (50, 51). Future studies should consider the possibility that changes in the subcellular localisation of RNA within mitochondria may be more transient.

Examining subcellular RNA localisation within mitochondria is technically challenging. The cell fractionation method coupled with RNase digestion is commonly used but can also result in false positives due to incomplete RNase digestion. A refinement of the fractionation and RNase treatment approach is to create mitoplasts via removal of the outer mitochondrial membrane, thus removing a significant source of potential contaminating nuclear-encoded transcripts prior to RNase treatment (22). Importantly however, we have demonstrated that our RNase treated isolated mitochondria have a similar degree of purity as mitoplasts (39). Other approaches utilising proximity labelling techniques such as APEX-Seq and Proximity-CLIP avoid these issues but require genetic modifications to cells and have so far only been suitable to a small number of *in vitro* models. Fluorescent in situ hybridisation (FISH) is a well-accepted approach to visualise RNA transcripts co-localised to mitochondria but is not a high-throughput technique, since it requires the design of probes specific to each RNA target. Jeandard et al. (36) recently developed a novel CoLoc-seq that creates transcript specific RNase digestion kinetics. When combined with a mock CoLoc-seq protocol, the process can identify RNase-resistant contaminants in the mitochondria and other organelles. Jeandard et al. (36) used this approach alongside single-molecule fluorescence in situ hybridisation (smFISH) to suggest that although there were a very small number of bona fide nuclear-encoded RNA transcripts in the mitochondria, their abundance was low and many that have previously been detected were most likely extra mitochondrial, including 5S rRNA and RMRP. The rarity of nuclear-encoded RNAs localised in the mitochondria is consistent with the work of Fazel et al. (52) who used a proximity labelling approach to suggest most are localised on the outer membrane. In the present study, our analytical approach (correlation with *Mt-co1*) allowed us to account for the possibility of artefacts due to residual transcripts surviving the RNase digestion and purification procedure. Indeed, this analysis correctly predicted the expected strong positive correlation with all mtDNA-encoded genes, and the negative correlation with the overwhelming majority of nuclear-encoded genes expected to be localised to the cytosol or nucleus.

Understanding the subcellular distribution of RNA has implications for the development of RNA-based therapies. Notably, there is increasing interest in targeting RNA-based therapeutic agents to the mitochondria to treat a range of metabolic and neurological diseases (53). Thus, characterising the effects of potent physiological stimuli known to induce mitochondrial adaptations (such as exercise) on RNA subcellular localisation not only provides important fundamental biological knowledge, but may also inform novel therapeutic strategies.

In summary, we report that rat skeletal muscle mitochondria contain a select population of small and long nuclear-encoded RNAs. The expression levels of several miRNAs were changed in the mitochondria of endurance trained when compared to sedentary rats. Further research is now required to investigate the nature of their presence in the mitochondrial subcellular environment and to elucidate their potential role(s) in the regulation of exercise training adaptations.

## METHODS

### Ethical Approval

All experimental procedures were approved by the Deakin University Animal Ethics Committee (G02-2019). All animals were housed and treated in accordance with standards set by Deakin University’s Animal Welfare Committee, which complies with the ethical and governing principles outlined in the Australian code for the care and use of animals for scientific purposes.

### Animal procedures

Wistar Kyoto 4-5 week old male rats (n=32) were obtained from the Animal Resource Centre, Perth, Western Australia. Rats were housed in pairs and maintained for a 1-week familiarisation period with a 12-hour light/dark cycle, constant temperature of 21 ± 2°C, and humidity levels between 40 and 70%. Rats had *ad libitum* access to standard chow diet and tap water. After a week of acclimatisation, 5-week old rats were randomly allocated into two groups: sedentary (Sed, *n*=16) and exercise trained (ExT, *n*=16).

### Exercise training

At 5-weeks of age, animals began the exercise training protocol which consisted of running 5 days per week for 4 weeks on a motorized treadmill (Exer 3/6, Columbus Instruments, OH, USA), as per our previous studies (37, 54, 55). Briefly, during week 1 of training, the running duration was progressively increased in 10-minute increments each day, from 20 up to 60 minutes, with speed set at 15 m/min. In weeks 2, 3 and 4, animals ran for 60 minutes at 20 m/min. Animals in the sedentary control group were placed on a stationary treadmill for the same duration while the exercise training group rats were running.

### Humane killing and tissue collection

At 9-weeks of age and 24 hours following the competition of the final session of the 4-week training protocol, animals were heavily anaesthetised (determined by loss of pedal reflex) with 5% isoflurane gas then humanely killed via exsanguination following removal of the heart. The heart was weighed then left ventricle wall was dissected and weighed. A technical fault led to inaccurate measurement of heart tissue mass for *n*=2 Sed and *n*=4 ExT animals; therefore, these data were excluded from analysis. A portion of left ventricle was separated for analyses on fresh tissue and the remainder was snap frozen. Gastrocnemius skeletal muscle was excised then the inner red portion was rapidly dissected and ∼100 mg pieces were used for mitochondrial isolation preparations (2-4 replicates per tissue). Approximately 10 mg of fresh tissue was used for analyses of mitochondrial respiration (see below), then the remaining tissue was snap frozen in liquid nitrogen for subsequent RNA and protein analyses.

### Mitochondrial respiration analysis

Upon dissection, tissue (left ventricle and red gastrocnemius muscle, ∼10 mg) was immediately placed in ice-cold BIOPS preservation media (7.23 mM K_2_EGTA, 2.77 mM CaK_2_EGTA, 5.77 mM Na_2_ATP, 6.56 mM MgCl_2_·6H_2_O, 20 mM taurine, 15 mM phosphocreatine, 20 mM imidazole, 0.5 mM dithiothreitol, and 50 mM K-MES at pH 7.1) for mitochondrial respiration analyses that was performed within 48 hours. Muscle fibres were gently mechanically separated using fine-tipped forceps then placed in BIOPS containing 50 mg/mL saponin for 30 min at 4℃ with agitation for permeabilisation of the sarcolemma. Permeabilised muscle fibre bundles were then equilibrated with ice-cold MiR05 respiration media (0.5 mM EGTA, 10 mM KH_2_PO_4_, 3 mM MgCl_2_·6H_2_O, 60 mM lactobionic acid, 20 mM taurine, 20 mM HEPES, 110 mM D-sucrose, and 1 mg/mL bovine serum albumin at pH 7.1). Two portions of the fibre bundles were blotted on filter paper for 5 s, and wet-weight mass was recorded using a microbalance (2 – 3 mg per replicate). Permeabilized muscle fiber bundles were assessed in duplicate using a high resolution respirometer (Oxygraph O2k; Oroboros Instruments, Innsbruck, Austria) in respiration buffer MiR05 at 37°C with O_2_ concentration maintained between 300 and 500 µM, similar to our previous work (56, 57). Briefly, a substrate, uncoupler, inhibitor titration (SUIT) protocol was performed. Malate (2 mM) and glutamate (10 mM) were first titrated to assess O_2_ flux due to mitochondrial complex I leak (LEAK CI). Oxidative phosphorylation (state 3 respiration) supported by CI substrates (OXPHOS CI) was then determined with the addition of ADP 2.5 mM. Cytochrome c (10 µM) was added to confirm inner mitochondrial membrane integrity which was accepted as being ≤15% increase in O_2_ flux. This was followed by succinate (10 mM) to assess oxidative phosphorylation (state 3 respiration) supported by convergent complex I & II substrate input (OXPHOS CI+II). Uncoupled respiratory capacity of the electron transfer system supported by convergent complex I & II substrate input (ETS CI+II) was determined after 2–4 titrations (1 - 2 µM) of carbonyl cyanide p-trifloromethoxyphenylhydrazone (FCCP). Inhibitors of specific complexes were then applied: rotenone (0.5 µM) to inhibit CI resulting in ETS supported only by CII substrate flux (ETS CII), followed by the CIII inhibitor antimycin A (2.5 µM) to determine residual non-ETS O_2_ flux that was subtracted from values in all respiratory states. Oxygen flux values were obtained using the average of both chambers during steady state for each respiratory state. If one of the chambers did not reach steady-state flux, that value was excluded from the analysis of that respiratory state.

### Skeletal muscle enzymatic activity assays

Frozen red gastrocnemius tissue was crushed to a fine powder in liquid nitrogen. Approximately 15-20 mg of the crushed tissue was homogenised in freshly prepared buffer (50 μL per mg tissue; 50 mM Tris, pH 7.5, containing 1 mM EDTA, 10% v/v glycerol, 1% v/v Triton-X100, 50mM NaF, 5mM NaPP (Na4P2O7), 1 mM PMSF and 5 μL/mL Protease Inhibitor Cocktail (cat# P8340, Sigma Aldrich, Australia). The tissue lysate underwent two freeze-thaw cycles and was then centrifuged at 10,000xg for 1 min at 4°C. The supernatant was transferred to a new tube and used for protein quantification (54, 58). Protein concentration was determined spectrophotometrically (562 nm) using a bicinchoninic acid (BCA) protein assay (cat# 23225, Thermo Fisher Scientific, Australia) using bovine serum albumin (BSA) protein standards (range 0.05-2 μg/μL). Citrate synthase activity was measured spectrophotometrically by measuring the increase in absorbance per minute of 5,5-dithiobis-2-nitroben-zoate (DTNB). The plate read was performed at 25°C at 412 nm for 5 min (54, 58–60). 3-hydroxyacyl-CoA dehydrogenase (β-HAD) activity was measured spectrophotometrically by measuring the decrease in absorbance per minute of nicotinamide adenine dinucleotide (NADH). The plate read was performed at room temperature at 340 nm for 5 min (54, 60). Enzyme activities were normalised to protein content and expressed as µmol/min/g protein.

### Mitochondrial isolation

Freshly dissected red portion of gastrocnemius muscle tissue (∼80 – 100 mg) was immediately placed in 1 mL ice-cold lysis buffer containing Protease Inhibitor Cocktail (5 μL/mL) (cat# P8340, Sigma-Aldrich, Australia), then minced using fine scissors before mechanical homogenisation with an ice-cold Teflon-tipped glass dounce homogeniser (10 passes with rotation at 350 rpm). Wash buffer (9 mL) was added to the lysate and intact mitochondria were immunoprecipitated using 50 uL anti-TOMM22 conjugated magnetic beads (#130-096-946, Miltenyi Biotek, Australia) with gentle inversion for 1 hour at 4℃. Lysate was then flowed through a pre-separation filter and column mounted on a magnetic rack and washed 3 times before being removed from the magnetic rack for elution. Eluate was centrifuged at 13,000 x g for 2 minutes, supernatant removed, and the mitochondrial pellet resuspended in 50 uL storage buffer. To obtain only RNA transcripts that were localised within intact mitochondria, the mitochondrial pellet was treated with RNase-A (312 μg/mL, cat# 19101 Qiagen, Australia) for 1 hour at 37°C. RNase-A activity was then halted by the addition of proteinase-K (3 mAU/mL, cat# 19131 Qiagen, Australia) then centrifuged at 8000 x g for 10 min at 4℃. The supernatant was aspirated, and the mitochondrial pellet was washed twice with 100 μL ice-cold Storage Buffer and centrifuged at 13,000g for 2-minutes at 4°C between washes. Finally, the supernatant was aspirated and the highly purified RNase-treated mitochondrial pellet was resuspended in 100 uL ice-cold storage buffer and frozen at -80℃.

### Isolated mitochondria RNA extraction

Frozen mitochondria samples were thawed on ice in the presence of 5 volumes of TRI-Reagent Solution (Zymo Research #R2061, Integrated Sciences, Australia), pipette mixed and then centrifuged at 16,000 x g for 1 min. RNA was extracted from isolated mitochondria using the Zymo Direct-Zol Microprep kit (Zymo Research #R2060, Integrated Sciences, Australia) with on-column DNase-I digest as per the manufacturer’s instructions. Mitochondrial RNA was eluted in 12 μL nuclease-free water and quantified by gel electrophoresis using the Tapestation High Sensitivity RNA screentape and reagents (Agilent Technologies #5067-5579 and 5067-5580, Integrated Sciences, Australia).

### Tissue RNA extraction

For RNA extraction from red gastrocnemius muscle and left ventricle tissues, ∼15 mg whole tissue was placed in a liquid nitrogen pre-chilled cyrotube with a 5 mm RNase-free stainless steel bead. Tissue was mechanically disrupted with 2 cycles at 4m/s for 10 s (MP Biomedical FastPrep). Tri-reagent was added then RNA was extracted from the lysed sample (Zymo DirectZol RNA Miniprep kit #R2050, Integrated Sciences, Australia) with on-column DNase-I digest as per the manufacturer’s instructions. RNA was tested for purity on Nanodrop, concentration with Qubit HS RNA assay (Thermo Fisher, Australia) then RNA integrity number (RIN) was determined using the Tapestation with High Sensitivity RNA reagents as above. All RINs from whole tissue RNA were ≥7.0.

### Whole-tissue and isolated mitochondria total RNA sequencing

Skeletal muscle tissue total RNA (50 ng) and isolated mitochondria total RNA (10 ng) were converted to cDNA libraries (Zymo-Seq RiboFree Total RNA kit #R300, Integrated Sciences, Australia). Briefly, ribosomal RNA was depleted, then remaining transcripts were fragmented and converted to cDNA using random priming. After second strand synthesis, the ends of the cDNA were enzymatically repaired and Illumina-compatible sequencing adaptors were ligated. Library size (TapeStation, Agilent) and concentration (Qubit) was assessed prior to sequencing. RNA sequencing was performed on the Illumina NovaSeq 6000 platform (Deakin University Genomics Centre) with 151 bp paired end reads, generating at least 40 million reads per library. Left ventricle tissue total RNA (100 ng) from a subset of animals was converted to cDNA libraries (Illumina TruSeq Stranded Total RNA Ribo-Zero H/M/R kit) and sequenced on an Illumina NovaSeq 6000 platform (Macrogen Oceania), generating at least 50 million paired-end 151 bp reads per library. Reads from all libraries underwent quality check and adapter trimming with FastQC and Cutadapt then mapped to the rat genome build mRatBN7.2 with Ensembl v106 annotations with STAR aligner v2.7.1a then counted at the gene level in htseq-counts using the Galaxy Australia platform (61). Gene counts were normalised to library size (counts per million, CPM) and genes with >1 CPM in all samples were included for analysis of differential expression using Voom/Limma in Degust v4.1.5 (62). Libraries from two samples (1 animal in each group) were considered outliers based on principle component analysis and excluded from downstream analysis. Genes with a false discovery rate (FDR) <0.05 were considered differentially expressed. PCA plots were generated in iDEP (63). Pathway analyses for enriched Gene Ontology Biological Process terms was performed based on fold-change using GAGE in iDEP v0.96 (63) with filtered genes (>1 CPM) as background and pathway significance threshold of FDR<0.2. Analysis of RNA localisation in isolated mitochondria was performed by ranking the Pearson correlation coefficient (*r*) between read counts of each gene with the read counts of *Mt-co1* across all isolated mitochondria libraries (Sed and ExT groups combined). Data were visualised GraphPad Prism (v10.1, Boston, MA).

### Isolated mitochondria small RNA sequencing

Small RNA libraries were prepared from 17.8±15.9 ng total mitochondrial RNA (*n*=13 exercise, *n*=12 sedentary) using the NEBNext Multiplex Small RNA Library Prep Set (#E7560S, New England Biolabs, Australia) with modifications (39). Briefly, the 3’ adapter, reverse transcription primer and 5’ adapter were diluted 0.3x and applied in sequential reactions as per the manufacturer’s protocol. Following PCR amplification, the cDNA library was cleaned using the Monarch PCR & DNA Cleanup Kit (5 μg) (#T1030L, New England Biolabs) using the 7:1 ratio of binding buffer to sample and was eluted in 27.5 μL nuclease-free water. The eluate containing the purified cDNA construct was first quantified using the 1X dsDNA HS Assay Kit (#Q33231, Thermo Fisher Scientific, Australia) on a Qubit 4.0 Fluorometer. Fragment size distribution was then assessed on the Agilent 4200 Tapestation using the HS-D1000 screentape and reagents (#5067-5584 and 5067-5585, Agilent Technologies, Australia). A total of 25 libraries passed quality checks and contained the anticipated peak corresponding to adapter-ligated miRNAs (approximately 160 bp). An equimolar pool containing all uniquely indexed small RNA libraries was then loaded across two lanes on a 6% Novex TBE polyacrylamide gel (#EC6265BOX, Thermo Fisher Scientific) for size selection and purification of the miRNA region via gel excision, as described previously (39), and the gel-excised regions were resuspended in 12 µL TE buffer. Fragment size distribution was then assessed on Tapestation with HS-D1000 screentape and reagents.

Prior to RNA sequencing, the gel-excised fragments corresponding to the miRNA and 190bp regions (5 µL each, or 5 µL nuclease-free water for a no-template control) were re-amplified in triplicate in 100 µL reactions using 1x Q5 High-Fidelity Master Mix (cat# M0492S, NEBiolabs., Inc.) with 2 µL each P5 and P7 primers for Illumina, and the reaction volume brought to 100 µL with nuclease-free water. The PCR conditions consisted of 1 min at 95°C, and 6 cycles of 15 s at 95°C (denature), 15 s at 57°C (anneal), 15 s at 72 °C. The re-amplified miRNA and 190 bp regions were then assessed for fragment size distribution on Tapestation with HS-D1000 screentape and reagents. To purify the libraries from excess primers and adapter-dimer, the reamplified library pool was cleaned using a 0.8x and then 1.3x ratio of AMPure XP beads (cat# #A63881, Beckman Coulter, Inc., USA) to sample as per the manufacturer’s instructions. The reamplified library pools were then assessed for fragment size distribution using Tapestation with HS-D1000 screentape and reagents, in which a large proportion of adapter-dimer was still present. A second round of gel-excision as described above removed most of the adapter-dimer.

A 2 nM loading pool containing 50% reamplified miRNA region, 25% reamplified 190bp region and 25% Phi-X spike-in was sequenced on the Illumina Novaseq 6000 platform (Deakin Genomics Centre), generating 2 x 50bp single-end reads. Following sequencing, quality control and pre-processing of fastq files was performed using fastp (fastp v0.20.0) to remove the read1 adapter sequence (AGATCGGAAGAGCACACGTCTGAACTCCAGTCA) and filter poor quality reads. Next, reads were mapped using blast (v2.9.0+) with switch -task blastn and word length set to 19 nt to miRbase rat mature miRNAs (mature.fa; accessible from miRbase (v22.1) (42) downloaded March 12 2018). Raw read counts for each mature miRNA were counted using a short perl script. In addition, a small RNA quality score was generated for each library and is presented as the percentage of reads mapping exons (coding sequences) per 10,000 reads. Analysis of differential mitochondrial miRNA expression between the exercise trained and sedentary rats was performed using DEseq2 (v1.36) in RStudio (v4.2.1). MiRNAs were considered for differential expression in the current study if they were detected above a threshold of ≥100 normalised read counts in >80% of samples (irrespective of group) as previously used by our group for analysis of circulating miRNA micro array data (64). This was previously suggested as the minimum threshold required to detect physiologically relevant miRNA expression levels (65, 66) as miRNAs expressed below this threshold displayed little evidence of gene repression in the cytosol of monocyte cells (66). Experimentally validated miRNA-mRNA targets were analysed using the multimiR package (database version 2.3.0, Bioconductor v3.15) (230) in RStudio (v4.2.1).

### Statistical analysis

All statistical analyses other than RNA-sequencing bioinformatics were conducted using GraphPad Prism (v10.1, Boston MA). Data are presented as mean(SD) and analysed by unpaired t-test or two-way ANOVA with adjustment for multiple comparisons as described in the respective figure legend.

## Supporting information

Supplementary Dataset 1 - LV RNAseq, ExT vs Sed

Supplementary Dataset 2 - SM RNAseq, ExT vs Sed

Supplementary Dataset 3 - Isolated Mito vs SM RNAseq

Supplementary Dataset 4 - Isolated Mito RNA & miRs, ExT vs Sed

## Data Availability

RNA-sequencing datasets from this study have been deposited at the NCBI Gene Expression Omnibus for the whole-transcriptome (accession number GSE267763) and miRNAs (accession number GSE220557).

## Author contributions

Conceptualization (animal experiments): JLS, SLamon, GDW, AJT

Conceptualization (RNAseq experiments): JLS, SLamon, SLoke, LC, MZ, GDW, AJT

Investigation (animal experiments): JLS, GM, GDW, AJT

Investigation (RNAseq experiments): JLS, SLoke, AJT

Investigation (mitochondrial respiration): AJT

Formal analysis (animal experiments): JLS, GDW, AJT

Formal analysis (bioinformatics): JLS, LC, MZ, DSH, AJT

Visualization: JLS, AJT

Writing – Original Draft: JLS, GDW, AJT

Writing – Review & Editing: JLS SLamon SLoke GM LC MZ DH GDW AJT

## Acknowledgments

We acknowledge Amandi Gunawardena for assisting with animal experimental procedures. Séverine Lamon is supported by a Future Fellowship from the Australian Research Council (FT210100278).

