## Supplementary Figures for "Skeletal muscle mitochondria contain nuclear-encoded RNA species prior to and following adaptation to exercise training in rats"

**This PDF file includes:**

Figures S1 to S2
Legends for Datasets S1 to S4

**Other supporting materials for this manuscript include the following:**

Datasets S1 to S4

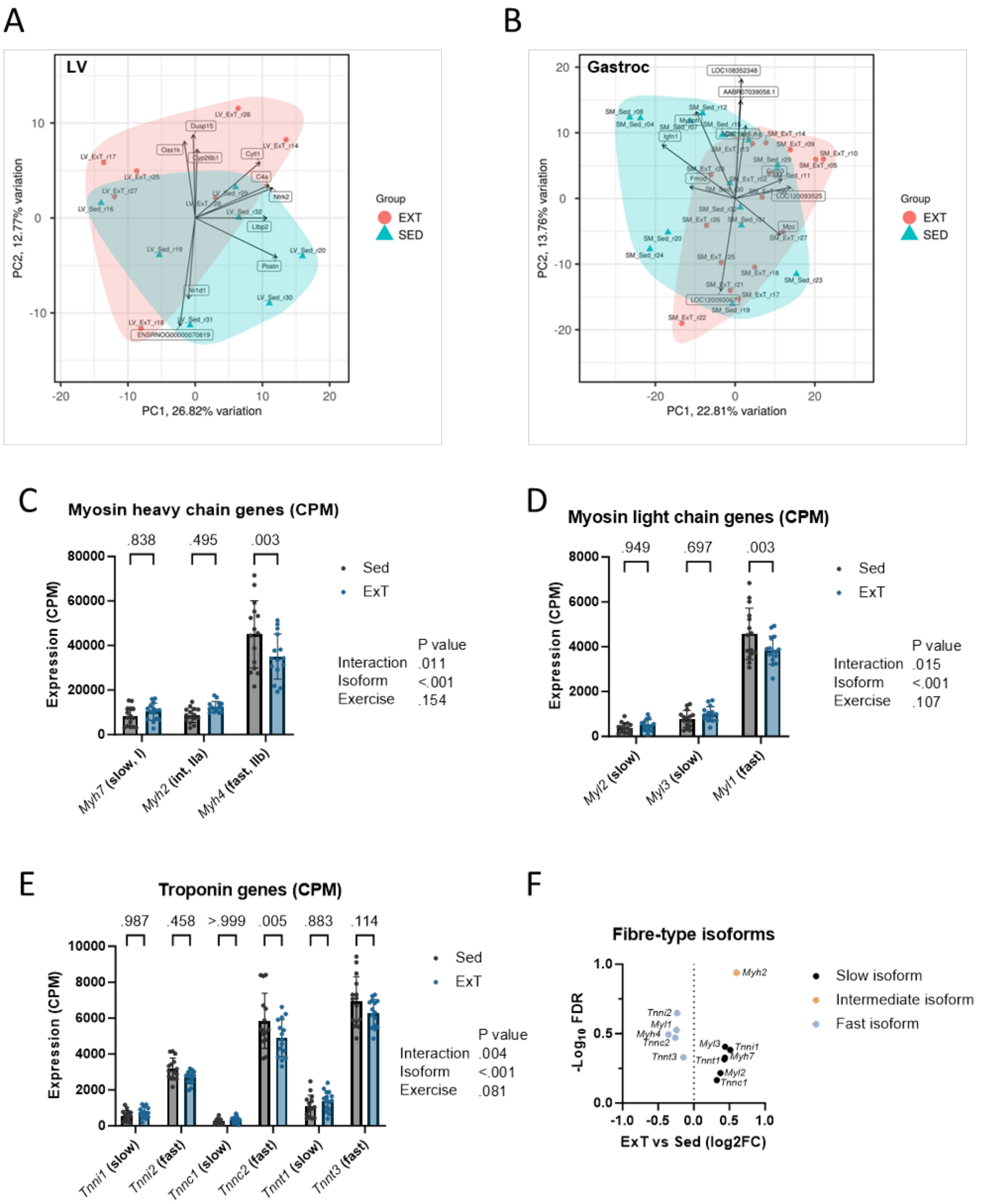


**Fig. S1.** Effects of exercise training on whole transcriptome and expression of muscle fibre-type genes. Principal component analysis of the whole transcriptomes of **A)** left ventricle tissue (*n*=7/grp) and **B)** gastrocnemius skeletal muscle tissue (*n*=15/grp). **C)** Effect of exercise trained (ExT) compared with sedentary control rats (Sed) on the expression of genes encoding isoforms of myosin heavy chain, **D)** myosin light chain and **E)** troponin known to be predominantly expressed in either slow type-I, intermediate type-Iia or fast type II-b rat skeletal muscle fibres. Data are normalised RNA-seq counts per million (CPM), mean(SD) for *n*=15/grp; two-way ANOVA with Šídák''s multiple comparisons test. **F)** Volcano plot summary of RNAseq analysis of myofibrilar related gene isoforms associated with slow, intermediate and fast isoform fibre-types, based on FDR adjusted RNAseq data.

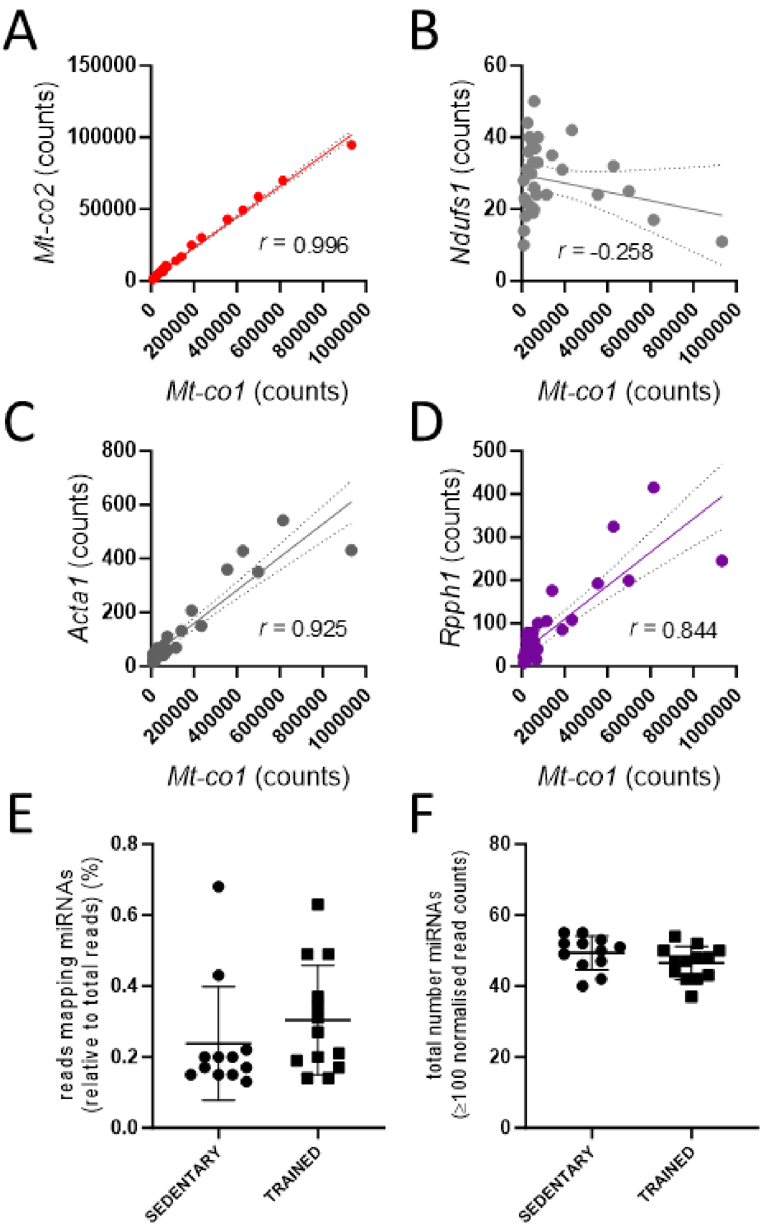


**Fig. S2.** Transcriptome of isolated mitochondria from red portion of rat gastrocnemius skeletal muscle. **A-D)** Correlation of representative genes with *Mt-co1* across all isolated mitochondrial samples (*n*=30 animals, Sed and ExT combined); red points represent mtDNA-encoded RNAs, grey and purple are nuclear-encoded mRNAs and ncRNAs, respectively. Positive correlation indicates mitochondrial localisation, negative correlation suggests non-mitochondrial localisation. **E)** Fraction of reads mapping to annotated miRNAs in mitochondrial samples. **F)** Total number of miRNAs detected per mitochondrial library with >100 read counts.

**Dataset S1 (separate file).** LV RNAseq, ExT vs Sed.xlsx. Normalised RNAseq count data, log2 fold-change and false discovery rate (FDR) values for left ventricle (LV) tissue from exercise trained (ExT) compared with sedentary (Sed) control rats.

**Dataset S2 (separate file).** SM RNAseq, ExT vs Sed.xlsx. Normalised RNAseq count data, log2 fold-change and false discovery rate (FDR) values for gastrocnemius skeletal muscle tissue from exercise trained (ExT) compared with sedentary (Sed) control rats.

**Dataset S3 (separate file).** Isolated Mito vs SM RNAseq.xlsx. Normalised RNAseq count data, log2 fold-change and false discovery rate (FDR) values for gastrocnemius skeletal muscle (SM) tissue and respective isolated mitochondria (MT) from exercise trained (ExT) compared with sedentary (Sed) control rats.

**Dataset S4 (separate file).** Isolated Mito RNA & miRs, ExT vs Sed.xlsx. **A)** whole transcriptome and **B)** microRNA normalised RNAseq count data, log2 fold-change and false discovery rate (FDR) values for isolated mitochondria (MT) from gastrocnemius skeletal muscle (SM) tissue in exercise trained (ExT) compared with sedentary (Sed) control rats.
